# NPD1/GPR37 axis protects painful traumatic brain injury and its complications

**DOI:** 10.1101/2024.05.20.594957

**Authors:** Junli Zhao, Sharat Chandra, Yuqing Wang, Vivian Zhang, Haichen Wang, Ru-Rong Ji

**Affiliations:** Center for Translational Pain Medicine, Department of Anesthesiology, Duke University Medical Center, Durham, NC 27710, USA; Department of Neurology, Duke University Medical Center, Durham, NC 27710, USA; Department of Cell Biology, Duke University Medical Center, Durham, NC 27710, USA; Department of Neurobiology, Duke University Medical Center, Durham, NC 27710, USA

## Abstract

Patients with traumatic brain injury (TBI) frequently exhibit heightened pain and associated complications such as cognitive decline, depression, and anxiety. GPR37 is widely expressed in various brain regions, but its function remains largely unclear. We recently discovered neuroprotectin D1 (NPD1) as a novel GPR37 ligand. In this study, we examined the protective role of the NPD1/GPR37 signaling pathway in TBI-induced neuropathic pain and its complications. TBI was induced by closed-head impact and resulted in transient neuropathic pain for less than two weeks, showing periorbital and cutaneous mechanical allodynia/hyperalgesia, as well as motor deficiency and cognitive impairment. We found that peri-surgical treatment with NPD1, effectively prevented TBI-induced mechanical hypersensitivity, motor deficiency, and cognitive impairment. NPD1 treatment also substantially inhibited TBI-induced microgliosis, astrogliosis (including A1 astrocyte markers), and neuroinflammation in the sensory cortex and hippocampus. RNA sequencing and GO enrichment analysis revealed downregulations of genes related to “calcium ion homeostasis,” and “GPCR signaling pathway” in the TBI-affected brain. These downregulations were restored by NPD1 treatment. RNAscope *in situ* hybridization revealed predominant *Gpr37* mRNA expression in oligodendrocytes. TBI resulted in rapid and remarkable demyelination and downregulation of *Gpr37* mRNA expression in oligodendrocytes, and both were protected by NPD1 treatment. NPD1’s inhibition of periorbital and cutaneous mechanical pain was abolished in *Gpr37^-/-^* mice. Moreover, TBI-induced neuropathic pain was prolonged by swimming stress, and NPD1 treatment prevented the stress-induced transition from acute to chronic pain in wild-type mice but not *Gpr37^-/-^* mice. Finally, chronic pain was associated with depression and anxiety, and NPD1 treatment mitigated these chronic pain complications through GPR37. Thus, through modulation of demyelination, diverse responses of glial cells, and neuroinflammation, the NPD1/GPR37 axis serves as a protective mechanism and a therapeutic target against painful traumatic brain injury and its complications.

## Introduction

Traumatic brain injury (TBI) is a significant neurological condition that affects brain function and a leading cause of mortality and long-term disability in the United States. According to the Centers for Disease Control and Prevention (CDC), TBI was responsible for approximately 190 American deaths daily in 2021, highlighting its severe public health impact. The CDC prioritizes the prevention of TBI and aims to minimize the likelihood of disability and other consequences for individuals who experience TBI. Individuals suffering from TBI often face chronic pain, which is a critical clinical concern due to its persistence and impact on quality of life. Chronic pain following TBI is complex, involving multifaceted interactions between neurological, psychological, and physiological processes(*1–6*). Clinical analyses of TBI revealed that patients frequently have elevated pain interference along with associated complications such as post-traumatic stress disorder (PTSD), depression, anxiety, and sleep disturbances(*7, 8*). Current efforts to prevent and treat acute and chronic pain induced by TBI have achieved limited success, in part due to our incomplete understanding of the molecular and cellular mechanisms underlying the pathogenesis of pain. Thus, there is urgency to identify the underlying mechanisms behind pain chronification and develop novel therapeutic approaches.

There is growing evidence that specialized pro-resolving mediators (SPMs), biosynthesized from fish oil DHA (docosahexaenoic acid), exhibit multiple benefits against inflammation and pain(*9, 10*). Especially, neuroprotectin D1 (NPD1), also called protectin D1 (PD1), have been shown to produce potent neuroprotective effects against ischemia brain injury, Alzheimer’s diseases, and neuropathic pain(*11–14*). Since NPD1 is biosynthesized from DHA, it emerges as a safer alternative that could potentially be integrated into dietary strategies for the treatment of acute and chronic pain. However, whether NPD1 prevents and reduces the development and severity of TBI induced acute and chronic pain and TBI-associated complications, as well as its underlying molecular and cellular mechanisms remains to be fully understood.

G-protein-coupled receptors (GPCRs) constitute the largest family of membrane receptors in humans and represent the largest family of drug targets of various therapeutics for neurological disorders including pain(*15, 16*). As a member of the orphan GPCR family, GPR37 is predominantly present in the brain, with expression noted in both neurons and glial cells(*17, 18*). GPR37 has attracted significant attention since its identification as a substrate of the E3 ubiquitin ligase parkin and its role in modulating dopaminergic neurotransmission in Parkinson’s disease(*19–21*). However, the role of GPR37 extends beyond the regulation of dopaminergic neurons. GPR37 is also recognized as the receptor for the neuroprotective and glioprotective factors prosaptide and prosaposin, and it serves as a key regulator of oligodendrocyte differentiation and myelination(*22–25*). Recently, we identified NPD1 as a novel ligand of GPR37, which mediates macrophage phagocytosis, promotes resolution of inflammatory pain, and confers protection against bacteria and parasite infections(*26, 27*). These studies indicate that GPR37 likely plays a more comprehensive role in regulating the functions of neuronal, glial, and immune cells. However, the function of GPR37 in TBI has not been investigated.

In this study, we found that *Gpr37* mRNA was highly expressed in oligodendrocytes in the sensory cortex and the hippocampus, two brain regions susceptible to TBI that are involved in the control of pain and cognition(*28*). Using a close-head injury model of TBI, our study demonstrated a powerful protective role of NPD1/GPR37 axis in neuropathic pain and its complications induced by TBI. Notably, NPD1 can effectively prevent the development of neuropathic pain and further treat established neuropathic pain in the TBI model through GPR37. Mechanistically, these beneficial effects of NPD1 stem from its powerful regulation of glial activation, neuroinflammation, and demyelination. Furthermore, NPD1 regulates its own receptor expression and homeostasis of cellular calcium and GPCR signaling in TBI animals.

## Results

### NPD1 prevents TBI-induced neuropathic pain, motor deficit, and cognitive impairment

To establish the model of pain induced by mild traumatic brain injury (TBI), we adopted a highly clinically relevant mouse model of mild closed head injury(*29*). In this TBI model, neuropathic pain was assessed by periorbital and plantar mechanical hypersensitivity on day 1, 3, 5, 7, 10 and 14 (**Fig. 1A**) in both sexes. Compared to Sham mice, TBI mice exhibited mechanical hypersensitivity starting from day 1 (*P* < 0.01, post-TBI, persisting for 10 days (*P* < 0.01) and full recovery by day 14, at both the periorbital site (primary) and plantar site (secondary) (**Fig. 1, B to E**). To interrogate the effect of NPD1 in the TBI model, we implemented treatments with NPD1. Specifically, NPD1 (500 ng) was administrated by intraperitoneal (i.p.) injections at 15 minutes before and 15 minutes after the TBI surgery. The efficacy of NPD1 was evaluated by testing mechanical allodynia and hyperalgesia at both the periorbital and plantar sites using von Frey methods.

**Fig 1.**
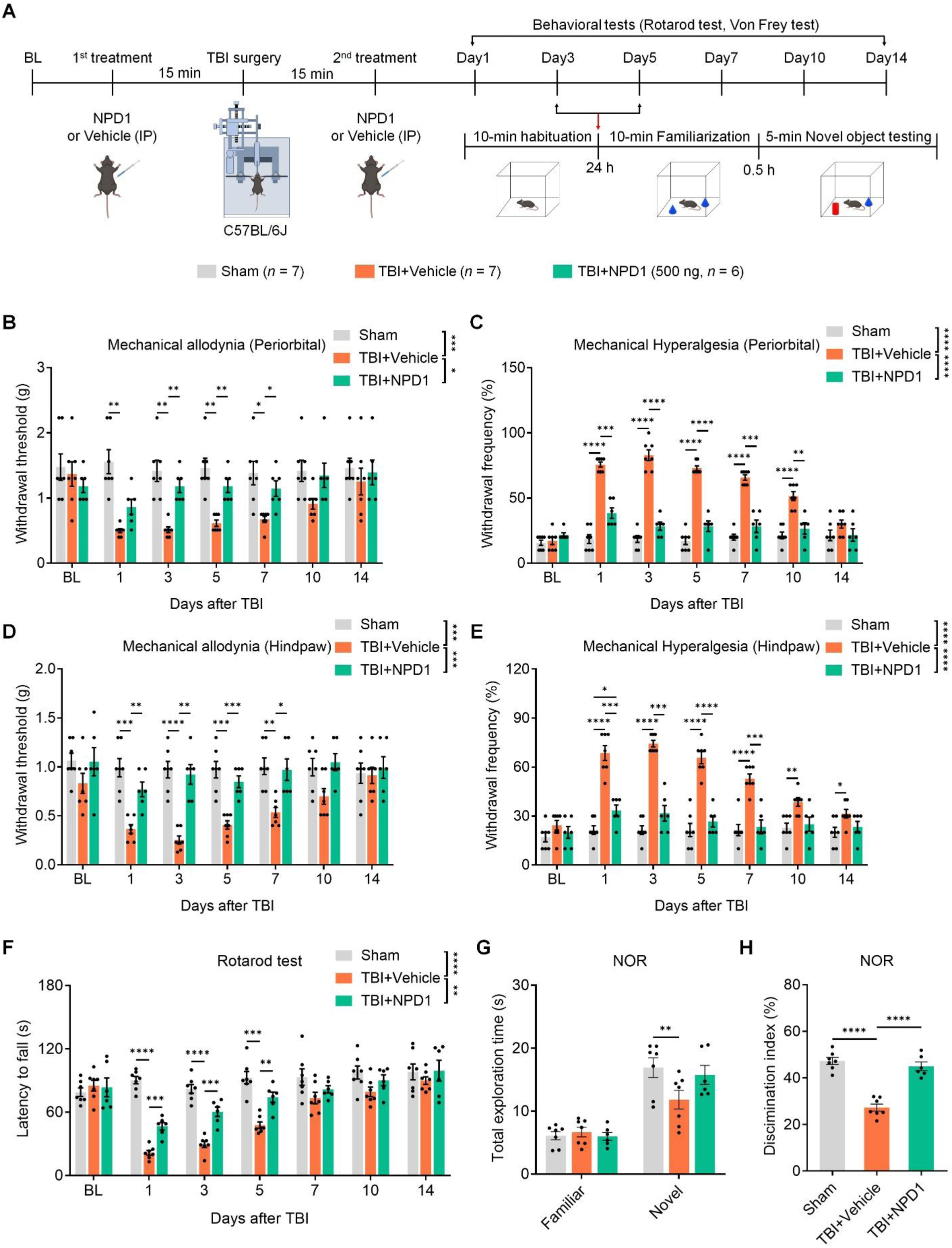
NPD1 prevents TBI-induced neuropathic pain and motor and cognitive impairment. (**A**) Experimental paradigm for close-head TBI, drug delivery, and behavioral test in WT mice with TBI or sham surgery. TBI mice were treated with vehicle, NPD1 (500 ng) by intraperitoneal (i.p.) injections, given 15 min prior to TBI and then 15 min after TBI. **(B, C)** Primary mechanical pain: von Frey testing showing withdrawal threshold (B) and withdrawal frequency (C) at periorbital sites in mice with sham surgery and TBI. **(D, E)** Secondary mechanical pain: von Frey testing showing withdrawal threshold (D) and withdrawal frequency (E) at hindpaw sites in the sham mice and TBI mice. **(F)** Motor coordination of sham mice and TBI mice in rotarod test, measured as latency to fall. **(G, H)** NOR test for learning and memory, showing total exploration time (G) and discrimination index (H) of sham mice and TBI mice. Data are represented as mean ± SEM. \**P* < 0.05, \*\**P* < 0.01, \*\*\**P* < 0.001, \*\*\*\**P* < 0.0001. Two-way ANOVA testing for two group comparison (B, C, D, E, F), Two-way ANOVA followed by Tukey’s multiple comparisons post hoc test for time point comparison (B, C, D, E, F). One-way ANOVA followed by Tukey’s post hoc test (G, H).

First, we measured primary orofacial pain in the periorbital region over the rostral portion of the eye (*i.e*., the area of the periorbital region facing the sphenoidal rostrum) of the mice. Compared to vehicle treated mice, NPD1 prevented the development of TBI-induced orofacial mechanical allodynia from day 3 (*P* < 0.01, **Fig. 1B**) and hyperalgesia from day 1 (*P* < 0.001, **Fig. 1C**). Next, we further measured secondary cutaneous pain at plantar sites. TBI mice with vehicle treatment exhibited robust mechanical plantar allodynia (**Fig. 1D**) and hyperalgesia (**Fig. 1E**), which were prevented by NPD1 treatment from day1 post TBI surgery. Overall, these data demonstrate that treatment of NPD1 is able to prevent TBI-induced pain hypersensitivity at periorbital and plantar sites.

Patients with mild TBI commonly present with balance disturbance, poor coordination, and acute cognitive impairment(*30*). We also tested motor coordination to determine the time course of motor recovery after TBI (**Fig. 1A**). Rotarod testing revealed that TBI mice exhibited transient motor coordination impairment from day 1 to day 5 after the injury, with noticeable recovery observed by day 7 (**Fig. 1F**). We found that NPD1 treatment significantly improved the motor function of TBI mice (**Fig. 1F**).

We also assessed the cognitive function of TBI mice using the novel object recognition (NOR) test to compare the discrimination index during days 3-5 post-injury (**Fig. 1A**). Compared with Sham mice, TBI mice with vehicle treatment spent less time exploring the novel object and exhibited a lower discrimination index (Sham vs. TBI + Vehicle = 47.26% vs. 27.26%) at 0.5 h (**Fig. 1, G and H**), indicating impaired cognitive function in this model. Notably, NPD1 treatment reversed the cognitive impairment in TBI mice, elevating the discrimination index from 27.26% to 44.98% (**Fig. 1H**). Therefore, NPD1 treatment effectively prevents not only neuropathic pain but also motor and cognitive impairments in mice with TBI.

### NPD1 prevents TBI-induced demyelination, gliosis, and neuroinflammation

Mild closed head TBI caused by rapid impact-acceleration-deceleration forces often lead to damages in the brain’s white matter, which primarily affects long axons that are essential for brain connectivity. When the intact axons are damaged, the myelin sheaths remain for prolonged periods, which may activate neuroinflammation. To determine whether our TBI model can cause demyelination, we tested the myelin in the cortex and hippocampus. We performed immunofluorescence (IF) analysis using antibodies against Myelin Basic Protein (MBP), the most common cellular marker for myelin (**Fig. 2A**). We found significantly decreased MBP levels in the cortex (*P* < 0.0001, **Fig. 2B**) and hippocampus (*P* < 0.0001, **Fig. 2C**) in mice subjected to TBI compared to mice with sham surgery. Further analysis confirmed that NPD1 treatment had a noticeable remyelinating effect, as shown by increased MBP IF, both in the cortex (*P* < 0.05, **Fig. 2B**) and hippocampus (*P* < 0.01, **Fig. 2C**).

**Fig. 2.**
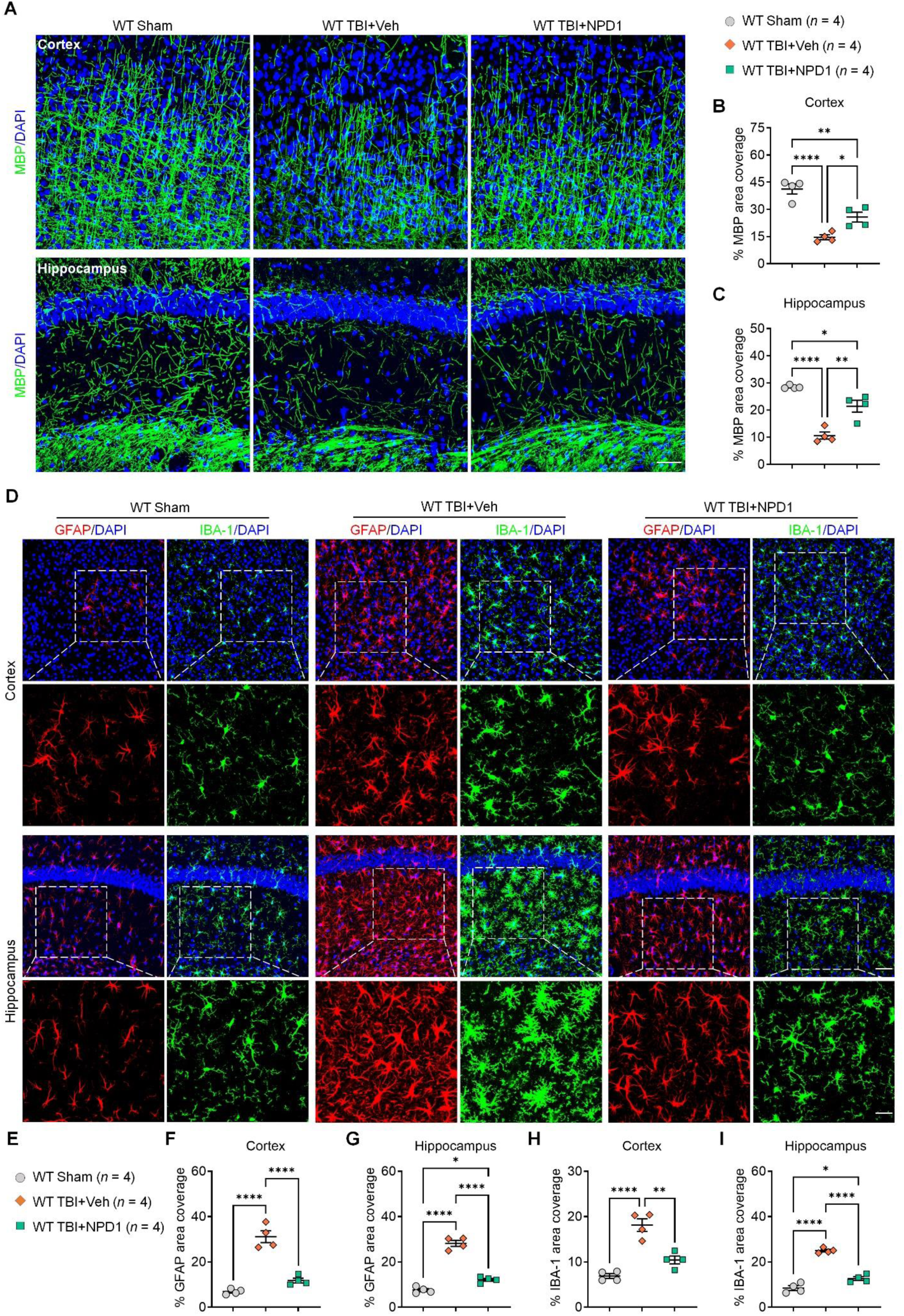
NPD1 pretreatment prevents TBI-induced demyelination and microglial and astroglial reactions. (**A**) Representative image of MBP immunostaining in the cortex and hippocampus in mice with sham surgery (Group-1) and TBI mice treated with vehicle (Group-2) and NPD1 (Group-3). Scale bar, 50 µm. **(B, C)** Analysis for MBP immunostaining as percent area covered by positive staining in the sensory cortex (B) and hippocampus (C). **(D)** Representative images of GFAP and IBA1 immunostaining in mice with sham surgery and TBI (treated with vehicle and NPD1) in the cortex (1^st^ row) and hippocampus (3^rd^ row). Scale bar, 25 µm. Inserted boxes are higher magnification images showing astrocytes and microglia in the cortex (2^nd^ row) and hippocampus (4^th^ row), scale bar, 50 µm. **(E-I)** Quantification of glial reaction. **E** Schematic of treatment groups and sample sizes for glial marker immunostaining. **F, G** Analysis for percent area covered by GFAP immunostaining in the cortical (F) and hippocampal (G) slices. **H, I** Analysis for percent area covered of IBA1 immunostaining in the cortical (H) and hippocampal (I) slices. Data are represented as mean ± SEM. \**P* < 0.05, \*\**P* < 0.01, \*\*\*\**P* < 0.0001. One-way ANOVA followed by Tukey’s post hoc test (B, C, F, G, H, I).

Next, we aimed to determine whether gliosis and neuroinflammation(*6*) are involved in the development of functional deficits of TBI and the protective effects of NPD1. To this end, we utilized IF to visualize astrocytes and microglia (**Fig. 2D**). We found significant astrogliosis in TBI mice treated with vehicle relative to mice with sham surgery in the cortex and hippocampus (*P* < 0.0001; **Fig. 2, D to G**). Interestingly, when TBI mice were treated with NPD1, there was a notable prevention of astrogliosis. This was evidenced by the quantitative IF analysis of GFAP in both the cortex (*P* < 0.0001, **Fig. 2F**) and hippocampus (*P* < 0.0001; **Fig. 2G**). In addition, to examine the effects of NPD1 on microglia reactivity (microgliosis), we preformed quantitative IF analysis with IBA-1, revealing an increase in TBI mice treated with vehicle relative to Sham mice that is significantly reduced in TBI mice treated with NPD1 in both the cortex (*P* < 0.01, **Fig. 2, D and H**) and hippocampus (*P* < 0.0001, **Fig. 2, D and I**).

To examine gene expression and function alteration in the TBI model, we performed Poly(A) RNA sequencing (RNA-seq) and gene differential expression analyses on cortical and hippocampal tissues from three groups of animals, 1) Sham surgery, 2) TBI surgery with vehicle treatment, and 3) TBI surgery with NPD1 treatment (**Fig. 3A**). Principal component analysis (**Fig. 3B**) demonstrated different groups as the main source of variation (40.9%). In RNAseq experiment, A total of 21,414 genes were analyzed, and a list of differentially expressed genes was included for various group comparisons. Volcano plot comparisons showed that TBI resulted in profound gene expression changes compared to the Sham group (Upregulated: 1128 and Downregulated: 1828) (**Fig. 3C**). Among the 1128 upregulated genes by TBI, 897 genes could be downregulated by NPD1 treatment. Furthermore, among the 1828 downregulated genes by TBI, 1414 were restored by the NPD1 treatment (**Fig. 3D, Fig. S1**). Compared to the Sham group, NPD1 treatment only produced very limited changes in gene expression (**Fig. 3E**), suggesting a powerful effect of NPD1 in preventing/restoring TBI-induced brain changes. Additionally, Gene Ontology (GO) enrichment analysis revealed that those genes that are upregulated by TBI but downregulated by NPD1 (897 genes) were implicated in pathways mainly related to “cytokine production,” “inflammatory response” and “immune response” (**Fig. 3F**).

**Fig. 3.**
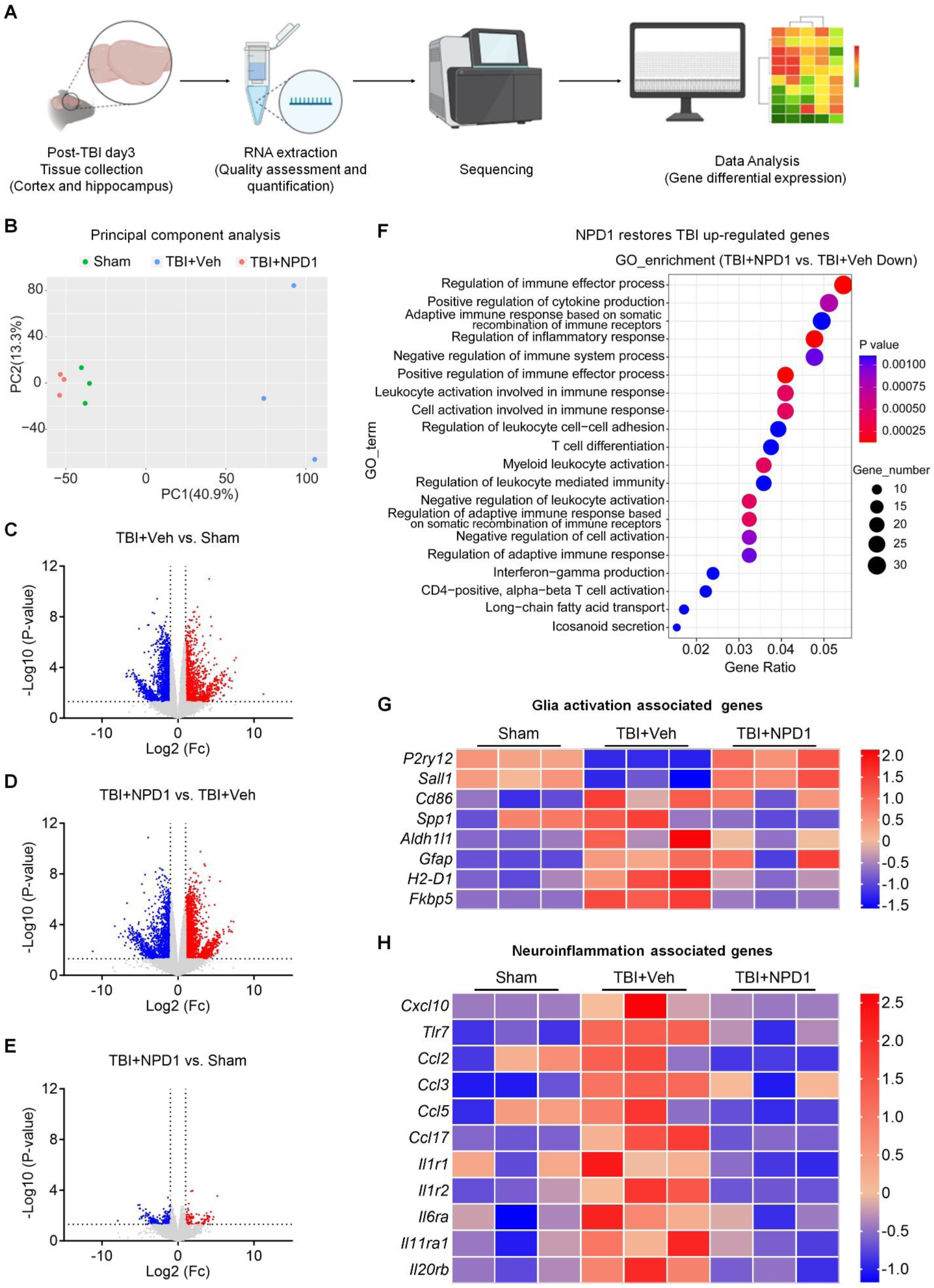
NPD1 confers protection against TBI-induced neuroinflammatory and glial responses. **(A)** Schematic of RNA-Seq in brain tissues from mice with sham surgery (Group-1) and TBI treated with vehicle (Group-2) or NPD1 (Group-3). **(B)** Principal component analysis showing sample clusterization by three treatment groups from 21414 variable genes. **(C-E)** Volcano plots of differential gene expression illustrating down-regulated (blue) and up-regulated (red) genes between Group-1 and Group-2(C), Group-2 and Group-3(D) and between Group-3 and Group-1 (E). Fold change > 2, *P* < 0.05. **(F)** GO enrichment analysis of up-regulated genes in Group-2 which are downregulated by NPD1 treatment in Group-3. **(G)** Heatmap depicting glia activation-associated genes that were altered in Group-2 but restored by NPD1 treatment in Group-3. **(H)** Heatmap depicting neuroinflammation-associated genes that were upregulated in Group-2, and restored by NPD1 treatment in Group-3. Data are represented as mean ± SEM. *n* = 3 animals for each group.

Next, we investigated glial reaction in the cortex and hippocampus after TBI and NPD1 treatment. We observed that the microglial “homeostatic” marker genes *P2ry12* and *Sall1* were downregulated, while the “activation” marker genes *Cd68* and *Spp1* were upregulated after TBI surgery, indicating marked microgliosis in this TBI model (**Fig. 3G**). Meanwhile, the astrocyte marker genes *Aldh1l1* and *Gfap* were upregulated after TBI surgery. Interestingly, the A1 reactive astrocyte specific genes *H2-D1* and *Fkbp5* were significantly upregulated in response to TBI surgery (**Fig. 3H**), highlighting the occurrence of astrogliosis. Our RNA-seq results showed that multiple pro-inflammatory genes, including *Cxcl10*, *Tlr7*, *Ccl2*, *Ccl3, Ccl5, Ccl17, Il1r1*, *Il1r2*, *Il6ra*, *Il11ra1* and *Il20rb* were upregulated in the TBI model (**Fig. 3H**). Notably, compared to the vehicle treatment, NPD1 treatment reversed TBI-induced gliosis and neuroinflammation (**Fig. 3, G and H**). These analyses demonstrate that NPD1 may prevent the progression of TBI-induced neuropathic pain and related complications by its control of glial reaction and neuroinflammation.

### NPD1 maintains GPCR signaling pathway and rescues *Gpr37* mRNA expression in TBI

GO enrichment analysis also revealed that TBI downregulated genes which were restored by NPD1 (1414 genes) were implicated in pathways related to “calcium ion homeostasis/concentration,” “GPCR signaling pathway” and “peptide receptor activity” (**Fig. 4A**). Since GPCR signaling pathway is known to mediate calcium dynamics and many biological functions(*31, 32*), NPD1 may achieve its biological actions in TBI through GPCR signaling. In our previous research, we established that NPD1 activates the GPR37 receptor in peripheral macrophages for the control of inflammation and inflammatory pain(*27*). Building on this, we extended our investigation into the brain’s NPD1-GPR37 interaction. We used 2D interaction mapping with 1000 ns simulation to demonstrate potential binding of NPD1 to GPR37 (**Fig. 4B**). Moreover, 3D molecular dynamics simulations confirmed that NPD1 forms hydrogen bonds with GPR37 at specific sites: TYR432, ARG418, GLU508, and GLN535(*27*) (**Fig. 4C**).

**Fig. 4.**
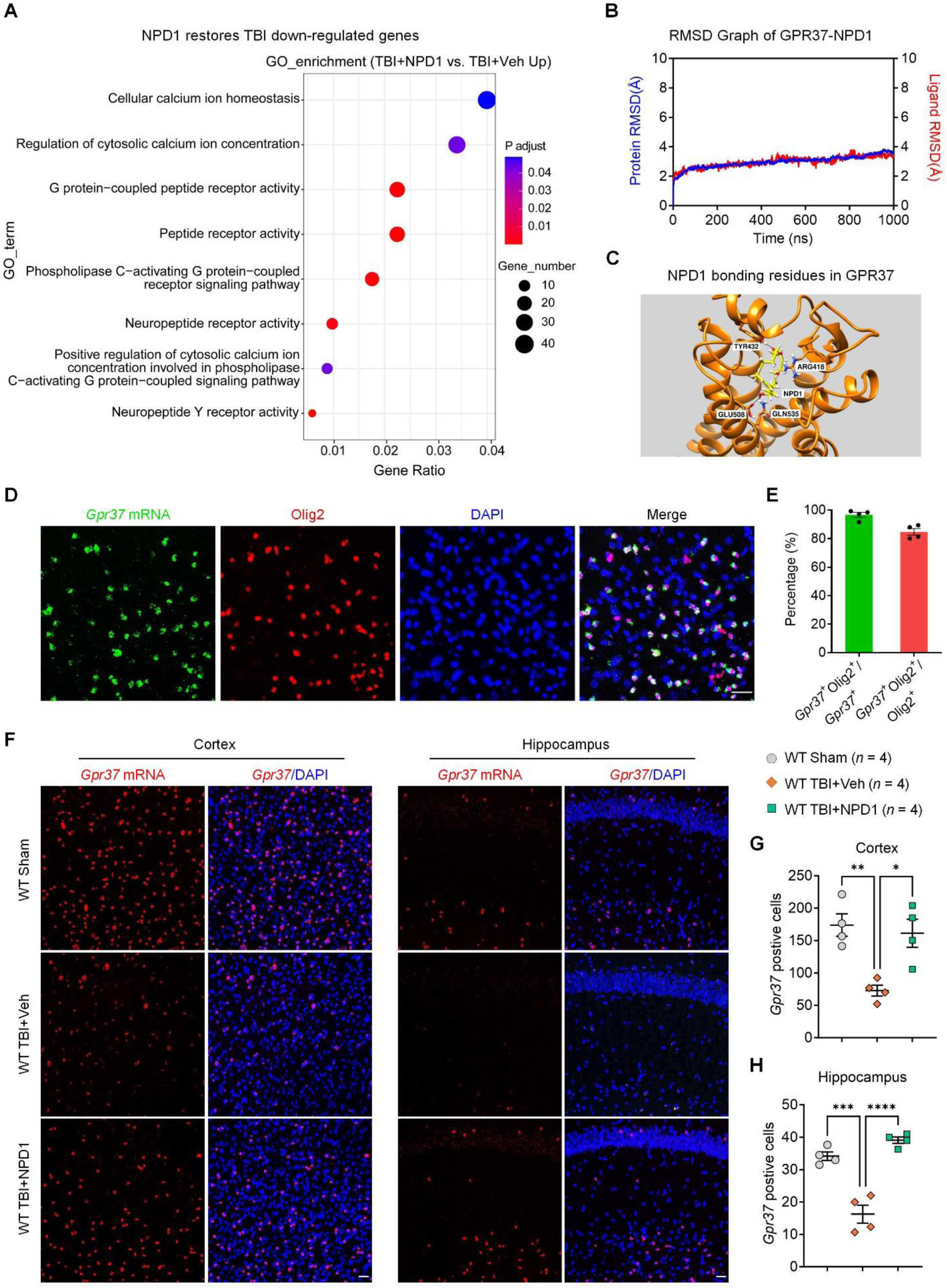
NPD1 protects against TBI-induced dysregulation of GPCR signaling pathway and downregulation of *Gpr37* expression. (**A**) GO enrichment analysis of down-regulated genes in TBI mice which could be reversed by NPD1 treatment. **(B)** Results of molecular dynamics simulation (MDS) for NPD1 interaction with GPR37. Y-axis shows root mean square deviation (RMSD graph) of the protein–ligand complex for 1000 nanoseconds (ns) time. The average 1000 ns RMSD graph was obtained by simulating the GPR37-NPD1 complex in three clusters for 1000 ns. The RMSD value of the GPR37 backbone (blue) and NPD1 atoms (red) are stabilized between 2 – 4 Angstrom. The overlapping RMSD line of both protein and ligand shows a stable binding in 1000 ns simulation. **(C)** Three-dimensional (3D) docking representation of GPR37-NPD1 interactions. NPD1 forms hydrogen bonds with the GPR37 residues TYR432, ARG418, GLU508, and GLN535. **(D)** Representative confocal images of double staining of *in situ* hybridization with RNAscope probes and immunofluorescence with Oligo2 antibody, showing co-expressing Olig2 and *Gpr37* mRNAs in cortical slices. Scale bar, 25 µm. **(E)** Quantification indicates that *Gpr37* is predominantly expressed by Olig2^+^ oligodendrocytes in the cortex. **(F)** Representative *in situ* hybridization images of *Gpr37* mRNA in WT mice with sham surgery (Group-1) and TBI treated with vehicle (Group-2) or NPD1 (Group-3) in the cortex and hippocampus. Scale bar, 25 µm. **(G, H)** Analysis of *Gpr37* mRNA positive cells in cortical (G) and hippocampal (H) slices from three groups. Data are represented as mean ± SEM. \**P* < 0.05, \*\**P* < 0.01, \*\*\**P* < 0.001, \*\*\*\**P* < 0.0001. One-way ANOVA followed by Tukey’s post hoc test (G, H).

To examine if NPD1 can activate GPR37 in our TBI model, we used *in situ* hybridization (RNAscope) to visualize *Gpr37* mRNA expression in the brain. Double staining revealed that *Gpr37* mRNA is highly co-localized with Olig2, the marker for oligodendrocytes (**Fig. 4, D and E**). Notably, *Gpr37* mRNA expression significantly decreased following TBI surgery, but NPD1 treatment was able to restore the *Gpr37* mRNA expression to normal levels (**Fig. 4, F to H**). These findings suggest that maintenance of GPCRs signaling and activation of GPR37 expression by NPD1 may play a protective role in the TBI model.

### NPD1 attenuates TBI-induced neuropathic pain through GPR37

To test the hypothesis that the NPD1’s protective action is dependent on GPR37, we evaluated the analgesic actions of NPD1 in both WT and *Gpr37^-/-^*mice on day 3 after TBI surgery (**Fig. 5A**). Initially, we employed the rotarod test to verify there is no motor impairment in all the treatment groups (**Fig. 5B**). On day 3 post-TBI, both WT TBI mice and *Gpr37^-/-^* TBI mice exhibited similar pain sensitivity at the periorbital and hindpaw sites (**Fig. 5, C to F**). Notably, a single injection of NPD1 (500 ng, i.p) on day 3 was able to mitigate TBI-induced mechanical allodynia and hyperalgesia at 1-, 2-, and 3-hours post-injection, with the effect diminishing after 24 hours, at both the periorbital and hindpaw sites. However, these antinociceptive effects of NPD1 were not observed in *Gpr37^-/-^*TBI mice (**Fig. 5, C to F**), demonstrating that the ability of NPD1 to reverse TBI-induced neuropathic pain through GPR37.

**Fig. 5.**
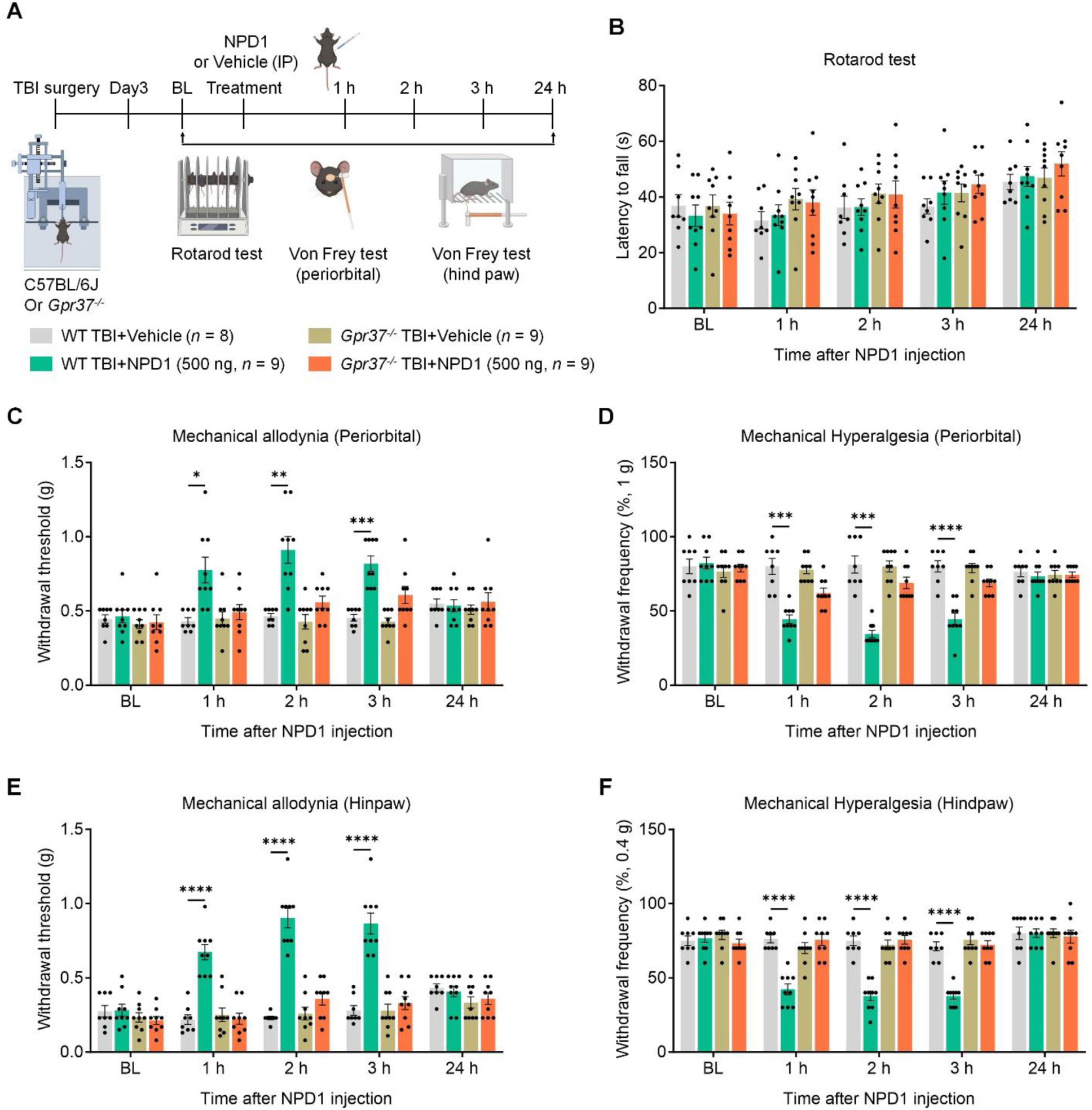
NPD1 post-treatment attenuates TBI-induced neuropathic pain through GPR37. (**A**) Schematic of experimental design for TBI surgery, drug delivery and behavioral test in WT mice treated with vehicle (Group-1) and NPD1 (Group-2) and *Gpr37^-/-^* mice treated with vehicle (Group-3) and NPD1 (Group-4). NPD1 (500 ng, i.p.) was given three days after TBI. **(B)** Rotarod test showing the motor function before and after NPD1 single injection. Note that NPD1 does not alter the motor coordination in the established TBI model. **(C, D)** Primary mechanical pain: von Frey testing showing withdrawal threshold (C) and withdrawal frequency (D) at periorbital site in WT and *Gpr37^-/-^* mice. **(E, F)** Secondary mechanical pain: von Frey testing showing withdrawal threshold (E) and withdrawal frequency (F) at hindpaw site in WT and *Gpr37^-/-^* mice. Data are represented as mean ± SEM. \**P* < 0.05, \*\**P* < 0.01, \*\*\**P* < 0.001, \*\*\*\**P* < 0.0001. Two-way ANOVA followed by Tukey’s multiple comparisons test for time point comparison (C, D, E, F).

### NPD1 prevents stress-induced development of chronic pain and the its complications in TBI animals via GPR37

Chronic pain is a complex condition influenced by various factors, especially by stress as a major risk factor(*33, 34*). To examine whether activation of the NPD1/GPR37 signaling pathway could prevent the development of chronic pain, we developed a model that combined TBI with swim stress, in which TBI was preceded by three days of forced swimming stress (SS) on days −3, −2, and −1 in WT mice and *Gpr37^-/-^* mice (**Fig. 6A**). In this combination model, we assessed periorbital and plantar mechanical hypersensitivity on days 3, 5, 7, 10, 14, 28, and 42 post-TBI surgery (**Fig. 6A**). While mechanical pain was fully recovered in TBI mice without stress (**Fig. 1, B to E**), stress substantially prolonged the duration of mechanical pain: there was no recovery even 42 days post-TBI in WT mice (**Fig. 6, B to E**). To evaluate whether NPD1 could prevent the development of chronic pain in this combination model, we administered a total of five injections of NPD1 (500 ng, i.p.) on days −3, −2, −1 (one hour before swimming stress), and then on day 0 (15 minutes before and after the TBI surgery) in both WT and *Gpr37^-/-^* mice. Strikingly, the NPD1 treatment fully prevented the stress-induced development of chronic pain at periorbital (**Fig. 6, B and C**) and plantar (**Fig. 6, D and E**) sites. This prevention began from day 3 (*P* < 0.05) and maintained through the entire time course of 42 days post-TBI (**Fig. 6, B to E**). However, the NPD1’s protection against chronic pain development was abolished in *Gpr37^-/-^* mice (**Fig. 6, B to E**). No sex differences were noted in the preventative effect of NPD1 in this model (**Fig. S2**). Collectively, these results indicate 1) stress is sufficient to cause a transition from acute pain to chronic pain after TBI; 2) NPD1 can prevent the development of stress-induced transition to chronic pain; and 3) GPR37 is crucial for the NPD1’s protection of chronic pain.

**Fig. 6.**
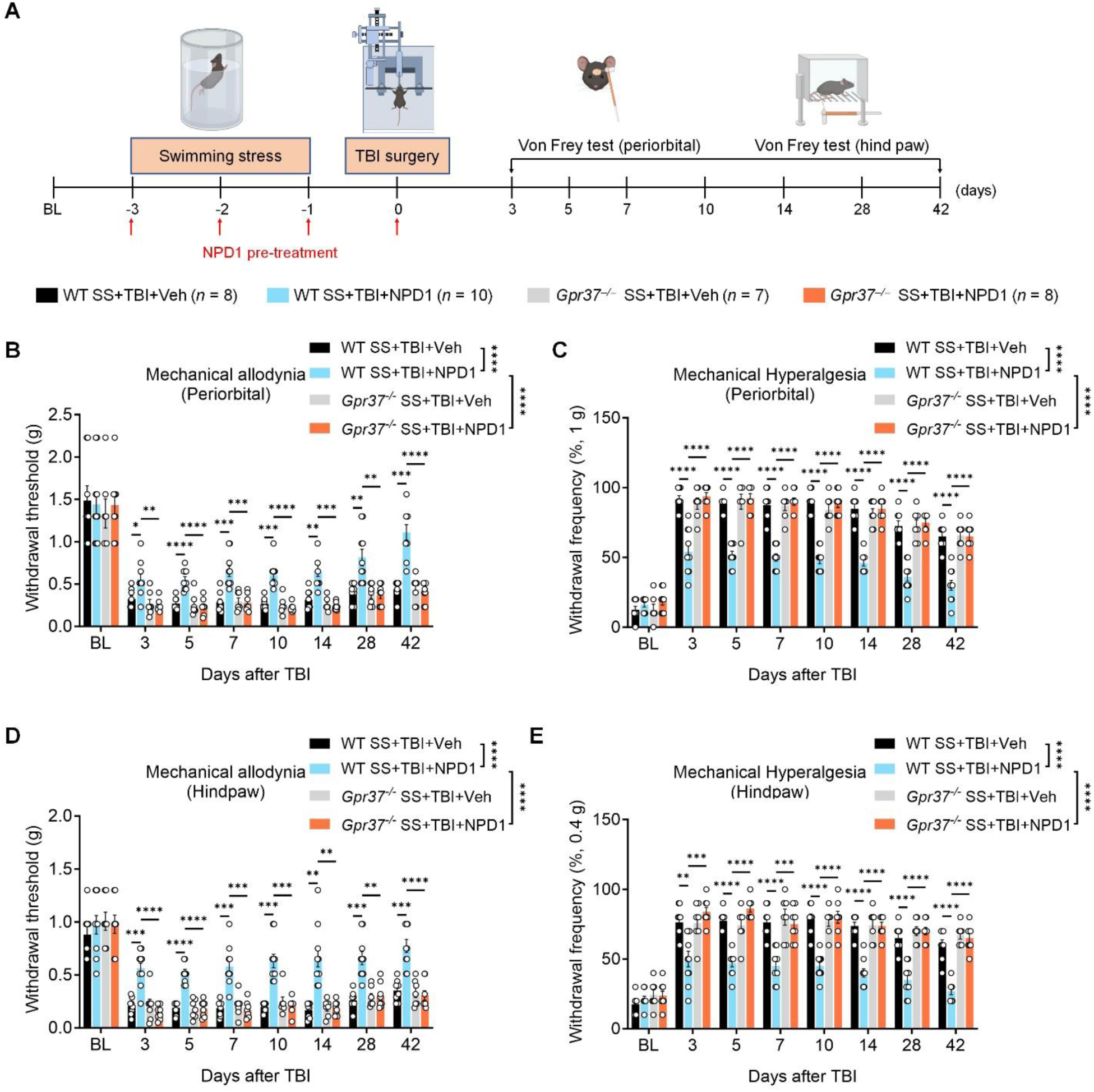
NPD1 prevents stress-induced chronification of pain through GPR37 in TBI mice with swimming stress. (**A**) Schematic of experimental design for drug delivery, swimming stress (SS), TBI surgery and behavioral test in WT mice with SS and TBI and treatment of vehicle (Group-1) and NPD1 (Group-2) and *Gpr37^-/-^* mice with SS and TBI and treatment with vehicle (Group-3) and NPD1 (Group-4). SS was induced daily for three days prior to TBI and NPD1 (500 ng, i.p.) was given daily prior to each SS. **(B, C)** Primary mechanical pain: von Frey testing showing withdrawal threshold (B) and withdrawal frequency (C) at periorbital site in four groups. **(D, E)** Secondary mechanical pain: von Frey testing showing withdrawal threshold (D) and withdrawal frequency (E) at hindpaw site in four groups. Data are represented as mean ± SEM. \**P* < 0.05, \*\**P* < 0.01, \*\*\**P* < 0.001, \*\*\*\**P* < 0.0001. Two-way ANOVA testing for two group comparison (B, C, D, E), Two-way ANOVA followed by Tukey’s multiple comparisons test for time point comparison (B, C, D, E, F).

Individuals with TBI often exhibit multiple post-concussion syndromes, including physical (e.g. poor motor coordination), cognitive (e.g. memory deficits), and emotional (e.g., anxiety and depression) comorbidities that significantly impact the quality of life. Stress can further exacerbate these detrimental effects(*34, 35*). Therefore, we not only assessed motor coordination function, but also investigated cognitive function using novel object recognition test, anxiety-like behaviors using the elevated plus maze and open field tests, and depression-like behaviors using the tail suspension test in the chronic pain model (**Fig. 7**). Moreover, we evaluated the beneficial effects of NPD1 treatment in these TBI-associated complications (**Fig. 7A**). In WT mice, NPD1 pretreatment significantly improved motor function in rotarod testing (**Fig. 7B**), reduced anxiety-like behaviors by increasing entries into the open arms in the elevated plus maze test (**Fig. 7, C to E**), and increasing entries into the center square in the open field test (**Fig. 7, F to I**), increased discrimination index in the novel object recognition (NOR) test at 0.5 hours and 24 hours (**Fig. 7, J to L**), compared to vehicle-treated mice. Furthermore, the NPD1 treatment group showed less severe depression-like behaviors, as indicated by reduced immobility in the tail suspension test in WT TBI mice (**Fig. 7M**). However, *Gpr37^-/-^* mice failed to demonstrate these beneficial effects of NPD1 (**Fig. 7, B to M**), suggesting that all the effects of NPD1 dependent on GPR37. Together, these data indicate that NPD1 pretreatment not only prevents the development of chronic pain but also mitigates the complications in this chronic pain model, highlighting a strong protective role of the NPD1/GPR37 signaling pathway against TBI-induced brain damage.

**Fig. 7.**
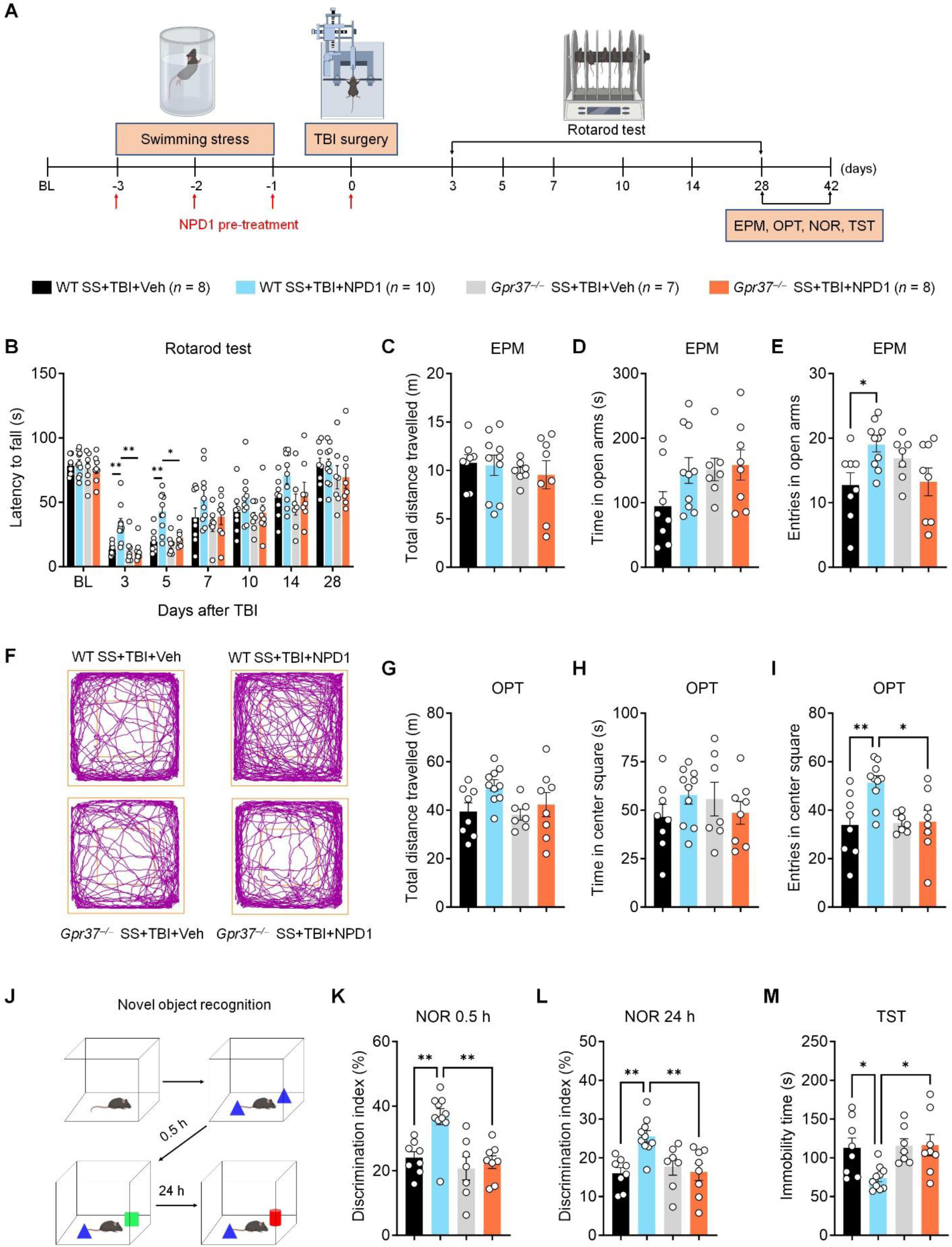
NPD1 prevents chronic pain related complications through GPR37 in TBI mice with swimming stress. **(A)** Schematic of experimental design for drug delivery, swimming stress (SS), TBI surgery and behavioral test in WT mice with SS and TBI and treatment of vehicle (Group-1) and NPD1 (Group-2) and *Gpr37^-/-^* mice with SS and TBI and treatment with vehicle (Group-3) and NPD1 (Group-4). SS was induced daily for three days prior to TBI and NPD1 (500 ng, i.p.) was given daily prior to each SS. **(B)** Motor coordination in rotarod test in four groups of WT and *Gpr37^-/-^* mice. **g** Schematic of NOR testing. (**C-E**) Elevated-plus maze test for anxiety-like behavior in four groups of WT and *Gpr37^-/-^*mice. **(F)** Representative travelled traces in open field test for four groups of WT and *Gpr37^-/-^* mice. **(G-I)** Open field test for anxiety-like behavior in four groups of WT and *Gpr37^-/-^* mice. **(J)** Schematic of NOR testing. **(K-L)** NOR testing showing discrimination index of four groups of WT and *Gpr37^-/-^* at 0.5 h (K) and 24 h (L). **(M)** Tail-suspension test for depressive-like behavior in four groups of WT and *Gpr37^-/-^* mice. Data are represented as mean ± SEM. \**P* < 0.05, \*\**P* < 0.01, \*\*\**P* < 0.001, \*\*\*\**P* < 0.0001. Two-way ANOVA followed by Tukey’s multiple comparisons test for time point comparison (B). One-way ANOVA followed by Tukey’s post hoc test (C, D, E, G, H, I, K, L M).

## Discussion

In this study, we have uncovered multiple protective roles of the NPD1/GPR37 signaling pathway in the damaged brain induced by TBI. First, we adapted a closed-head mild TBI model, which exhibited heightened acute pain sensitivity, impaired motor coordination, and cognitive deficiencies. Using this clinically relevant model, we discovered that peri-surgical NPD1 treatment (500 ng per mouse, i.p.), administered 15 minutes before and 15 minutes after TBI, effectively prevented sensory (neuropathic pain), motor, and cognitive disorders. Second, we observed robust glia responses, including oligodendrocyte-related demyelination and microgliosis and astrogliosis in the sensory cortex and hippocampus following TBI, which were effectively mitigated by NPD1. Consistently, NPD1 normalized the TBI-induced inflammatory responses and restored the altered gene expression for glia activation and neuroinflammation. Third, we found that NPD1 treatment resolved dysregulation of GPCR signaling caused by TBI. We also revealed interactions of NPD1 with GPR37 in the brain, as NPD1 upregulated *Gpr37* mRNA expression in the TBI-affected brain but lost its beneficial effects in *Gpr37^-/-^*mice with TBI. Lastly, we established that swimming stress caused acute to chronic pain transition following TBI. Using this stress-TBI combination model, we further verified that the NPD1/GPR37 signaling pathway confers prevention against the development of chronic pain. Importantly, the TBI-induced complications, including motor deficiency, anxiety and depression-like functional deficits, and cognitive impairment, which are associated with the chronic pain phenotype, were all mitigated by NPD1 treatment. Our data suggest that activation of the NPD1/GPR37 axis in the brain could effectively reduce neuroinflammation and glial responses and maintain the GPCR functions, thereby protecting against TBI-induced neuropathic pain and its complications.

TBI is known to cause prolonged neuroinflammation that can last for months or even years(*36–38*). Compared to open-head TBI models, our closed-head mild TBI model is more clinically relevant(*29, 38*). As previously reported(*39*), our TBI model displayed robust glia activation and inflammatory responses in the brain regions involved in sensory function and cognition. We observed significant activation and gliosis of astrocytes and microglia, evidenced by the marked increases in GFAP and IBA1 expression. The assessment of GFAP and IBA1 expression in the cortex and hippocampus revealed that NPD1 effectively prevented astrogliosis and microgliosis following TBI. Additionally, our RNAseq analysis validated gliosis by showing the upregulations of astrocyte and microglia activation associated genes, such as *Cd86*, *Spp1*, *Aldh1l1*, and *Gfap*. Specifically, P2ry12 is a microglia-specific marker in the healthy, unperturbed brain. When homeostasis is disrupted, microglia activation is characterized by both the downregulation of “homeostatic” markers (e.g., P2ry12, Fcrls, and Sall1) and the upregulation of “activation” markers (e.g., IBA1, CD68, CD206, CD45, and SPP1(*40*)). In our TBI model, both the downregulations of *P2ry12* and *Sall1* and upregulations of *Cd68* and *Spp1* were restored by NPD1 treatment, suggesting a powerful regulation of microglial phenotype by this pro-resolution mediator. Concurrently, the general astrocyte marker genes *Aldh1l1* and *Gfap* were also upregulated after TBI. Moreover, A1 reactive astrocytes produce pro-inflammatory substances and neurotoxins that can induce neuronal toxicity and cell death(*41*). The increase in A1-like astrocyte-specific marker genes *H2-D1* and *Fkbp5* following TBI surgery could therefore contribute to enhanced neuroinflammation in the brain. Additionally, both TBI and NPD1 also regulate neuroinflammation-related genes, including *Cxcl10*, *Tlr7*, *Ccl2*, *Ccl3, Ccl5, Ccl17, Il1r1*, *Il1r2*, *Il6ra*, *Il11ra1* and *Il20rb*. Thus, controlling neuroinflammation in the TBI model with NPD1 is a crucial mechanism for mitigating the development of neuropathic pain.

Myelin sheaths insulate nerves in both the peripheral and central nervous systems (CNS). Demyelination in the CNS can lead to various deficits, such as sensory and cognitive deficits(*42*). TBI was found to produce cerebral demyelination, mechanical damage to axons, and impairments in remyelination(*43*). Mild TBI results in significant and long-lasting demyelination in the white matter(*44*). We hypothesize that in our TBI model demyelination may trigger microglial and astroglial reaction and neuroinflammation (**Fig. S3)**. Both the initial injury and subsequent secondary processes can damage nerve fibers and disrupt the myelin sheath, resulting in demyelination, which can exacerbate the persistent pain and complications experienced by TBI patients. In the acute TBI model, we observed significant demyelination, as indicated by the marked decrease in MBP expression. The evaluation of MBP expression in the cortex and hippocampus showed that NPD1 effectively promoted remyelination following TBI surgery. Therefore, controlling demyelination via NPD1 signaling after TBI surgery is crucial for alleviating pain and related complications.

GPCRs regulate many critical brain functions in health and disease(*45*). In this study, we observed that the GPCR signaling pathway and related functions were downregulated by TBI, while NPD1 restored the changes. Although GPCRs are well investigated as therapeutic targets, more than 140 GPCRs remain mysterious and are referred to orphan GPCRs, which have gained considerable attention as drug targets recently(*46*). We uncovered that *Gpr37* mRNA is widely expressed in the brain, including the cortex and the hippocampus, two crucial regions affected by TBI. TBI led to a substantial decrease in *Gpr37* expression, which can be effectively prevented by NPD1. We also found that *Gpr37* mRNA is highly expressed in oligodendrocytes in the cortex and hippocampus (**Fig. S3)**. GPR37 was identified as a negative regulator of myelination, as *Gpr37^-/-^* mice exhibited hypermyelination of the corpus callosum(*22*). We found NPD1 can protect against TBI-induced GPR37 downregulation and demyelination in the brain. It remains to be investigated how NPD1 regulates GPR37-mediated myelination in different CNS regions.

In summary, the NPD1/GPR37 axis plays a significant role preventing the development of neuropathic pain and motor deficit in the closed-head TBI model. Furthermore, activation of the NPD1/GPR37 pathway prevented the onset and progression of chronic pain induced by stress combined with TBI. This combined model, which incorporates elements of physical trauma and stress, is particularly relevant to the clinical scenario as it closely mirrors the pain and complications observed in TBI and chronic pain patients experiencing stress and opioid addiction(*47, 48*). By activating GPR37 with NPD1, we can also effectively prevent the development of cognitive and emotional complications as chronic pain comorbidities. The implications of these findings are profound, as they open potential avenues for developing treatments that go beyond symptom management and address the root causes of pain and neurodegeneration following brain injury. Additionally, the fact that NPD1 is derived from DHA, a component found in fish oil, suggests the possibility of dietary interventions as part of a comprehensive treatment plan for TBI patients. Our findings underscore the importance of exploring the functions of GPR37 in the CNS and highlight the potential for novel therapeutic strategies that leverage the neuroprotective properties of NPD1 to treat the complex array of symptoms experienced by TBI patients.

## Materials and Methods

### Reagents

Neuroprotectin D1 (Item No. 10010390) was purchased from Cayman Chemical. Neuroprotectin D1 was dissolved in sterile normal saline and administered by intraperitoneal (i.p.) injection. The concentration of ethanol in the final solution was less than < 5%. The vehicle was sterile normal saline containing the same ethanol concentration.

### Animals

Adult mice (8-16 weeks old) of both sexes were used for behavioral tests. Wild-type (WT) mice (Stock No: 000664) and *Gpr37* knockout mice with a C57BL/6 background (Stock No: 005806) were purchased from Jackson Laboratory and maintained at Duke University Animal Facilities. All mouse procedures were approved by the Institutional Animal Care & Use Committee of Duke University. Mice were housed under a 12-hour light/dark cycle, with food and water available ad libitum. Animals were randomly assigned to each experimental group, with two to five mice housed per cage. All behavioral measurements were conducted during the daytime (light cycle). Animal experiments were conducted in accordance with the National Institutes of Health Guide for the Care and Use of Laboratory Animals. The experiments were performed and analyzed by researchers blinded to the genotype or treatment of the subjects.

### Mouse model of traumatic brain injury

The mild traumatic brain injury (TBI) model was adapted from our previously established model of closed-head injury(*29*) using WT or *Gpr37* knockout mice. After induction of anesthesia with 4.0% isoflurane, the trachea was intubated, and the mice were mechanically ventilated with 1.5% isoflurane in a 30% O_2_/70% N_2_ mixture. The mouse’s head was secured in a stereotactic device, and a midline scalp incision was made to identify anatomical landmarks. A concave 3-mm metallic disc was adhered to the skull, immediately caudal to the bregma. A single midline impact was delivered to the center of the disc surface using a 2.0-mm diameter pneumatic impactor (Air-Power Inc., High Point, NC), discharged at 6.5 ± 0.2 m/s, causing a head displacement of 2.2 mm. The anesthesia was discontinued after impact and the scalp incision was closed with suture. The animals were allowed to recover spontaneous respiration before extubation. Mice had free access to food and water post-surgery. Sham surgery was included as control, in which mice received identical surgical procedure without brain injury by impact.

### Swimming Stress

Swimming stress was performed daily for 3 days(*49*). On day 1, mice were placed in a glass container filled with 20 cm of water maintained at a temperature range of 22-24°C, for a duration of 10 minutes. On days 2 and 3, the duration was extended to 20 minutes. After each session, mice were gently dried with a towel and then placed in a drying cage with a heat lamp positioned above and a heat pad placed underneath, ensuring the mice were comfortably warmed and dried.

### Periorbital sensitivity testing for primary mechanical pain

Prior to baseline response assessment, mice were handled and habituated for three days. For both habituation and testing, mice were restrained in a cylindrical plastic tube with their heads exposed and secured by folding their tails over a napkin. The testing focused on the periorbital region above the rostral portion of the eye, near the area facing the sphenoidal rostrum. After a 5-min habituation period, von Frey filaments were applied perpendicularly to the periorbital skin in logarithmic force increments (Stoelting co., ranging from 0.008 to 2.0 g), starting from 0.16 g. Each filament was pressed with enough force to slightly buckle and was held for approximately 5 sec to induce a response. A positive response was identified by any of the following behaviors: vigorous face stroking with the forepaw, head withdrawal, or head shaking. In the absence of a response after 5 sec, a heavier filament was used. Conversely, a positive response prompted the use of a lighter filament. This process continued until six measurements were obtained for each mouse, or until four consecutive positive or negative responses were recorded. The 50% mechanical withdrawal threshold was then calculated. Periorbital hyperalgesia was assessed by measuring the response frequency to a higher force filament (1.0 g). This filament was applied 10 times to the periorbital site, each application lasting for 5 sec, with a 20-sec interstimulus interval between each application. The number of positive responses to the stimulus was recorded. The response frequency was then calculated using the formula: (number of positive responses / 10) × 100.

### Plantar sensitivity testing for secondary mechanical pain

Prior to the establishment of baseline responses, mice were handled and habituated over a three-day period. On both habituation and testing days, the mice were placed in plastic chambers set atop a wire mesh table, allowing access to their plantar sites. To assess mechanical allodynia, von Frey filaments ranging from 0.008 to 2.0 g (Stoelting Co.) were applied to the plantar surface of the hindpaw. The 50% withdrawal threshold was calculated using the up-down method. For measuring mechanical hyperalgesia, the response frequency was determined by applying a 0.4 g filament to the plantar site 10 times. The response frequency was then calculated using the formula: (number of withdrawals / 10) × 100.

### Rotarod test

Rotarod test (4-40 rpm accelerating mode) was conducted before TBI induction (baseline) and on days 1, 3, 5, 7, 10, and 14 after TBI. For mice subjected to both swimming stress and TBI surgery, the rotarod test was conducted before swimming stress (baseline) and on days 3, 5, 7, 10, 14, and 28 after TBI. After falling off the rotarod, the mice were returned to their home cages, and the latency to fall was automatically recorded by the rotarod system (IITC Life Science Inc.). If the mice remained on the rotarod for more than 300 sec, the latency was recorded as 300 sec.

### Elevated plus maze test

The elevated plus maze (EPM) apparatus (Stoelting Co., Item No. 60140) consists of two open arms (35 × 5 cm^2^) and two closed arms (35 × 5 × 15 cm^3^), elevated 50 cm above the ground. Immediately before the testing, mice were placed in the center of the maze, facing one of the open arms, and allowed to explore for 6 minutes. Anxiety-like behavior was assessed based on the time spent in the open arms and entries entered in the open arms. Both video and data were automatically recorded and analyzed using the ANY-maze software (Stoelting Co., Item No. 60000).

### Open filed test

Open field test (OPT) was conducted in a square arena measuring 45 × 45 × 45 cm^3^ (TAP Plastics) for a duration of 10 min. The center of the arena was designated as a central area encompassing 50% of the total arena space. Metrics such as the total distance traveled, time spent in the central area, and the number of entries into the central area were automatically recorded and analyzed using the ANY-maze software (Stoelting Co., Item No. 60000).

### Novel object recognition test

Novel object recognition (NOR) test was conducted as previously described(*50*). Mice were habituated on the first day by placing them in a 30 × 30 × 30 cm^3^ square arena (TAP Plastics). On the following day, two identical objects were positioned in distinct corners of the arena. After this exposure, the mice were returned to their home cages for a retention interval of either 0.5 hours or 24 hours. Subsequently, one of the two identical objects was replaced with a novel object for the recognition test. Post-retention, animals were reintroduced into the arena for a 5-min exploration period. Valid exploration was defined as the mouse either touching an object with its nose or closely focusing on it from a distance of less than 1 cm. Actions such as turning around, climbing, or sitting on the object were deemed invalid exploratory behaviors. Exploration times for the familiar and novel objects were evaluated using a discrimination index, calculated as DI = (T_N_ − T_F_)/(T_N_ + T_F_) × 100%. Animal behaviors were video-recorded and assessed by an experimenter blind to the test conditions.

### Tail suspension test

For tail suspension test (TST), mice were suspended by their tails using adhesive tape, positioned approximately 1 cm from the tip of the tail and about 15 cm above the surface. To prevent the animals from climbing or hanging onto their tails, plastic tubes were fitted over their tails. The test duration was set for 6 min, during which the mice were video recorded. A mouse was classified as immobile if it remained passive and motionless for a continuous period of at least 2 seconds. An experimenter, blind to the test conditions, quantified the duration of immobility.

### Brain collection and preparation for histochemistry

At the endpoint of the study, mice were terminally anesthetized using high concentration of isoflurane (∼10%) before transcardiac perfusion with PBS and 4% paraformaldehyde (PFA). This was followed by brain dissection and subsequent immersion in 4% PFA. After a 24-hour fixation period in PFA, the fixative was replaced with 30% sucrose, for an additional 24 hours. Brain sections, 30 μm thick, were then cut using a cryostat (Leica CM1950) and maintained as free-floating sections for immunostaining. Some sections were mounted onto the charged slides (Fisher scientific, Cat. No: 1255015) for *In situ* hybridization.

### *In situ* hybridization

*In situ hybridization* was performed using RNAscope® Multiplex Fluorescent Reagent Kit v2 (Advanced Cell Diagnostics, Cat# 323100) according to the manufacturer’s instructions. We used probe directed against mouse *Gpr37* (Advanced Cell Diagnostics, Cat# 319291). Following the completion of the RNAscope protocol, immunohischemistry was performed for double staining as described in the next section.

### Immunohischemistry

Sections were washed three times in PBS and blocked with 0.1% Triton X-100 and 10% goat serum for 1 hour at room temperature. The sections were then incubated overnight at 4°C in a humidified chamber with the following primary antibodies: anti-IBA-1 antibody (rabbit, 1:500, Wako, Cat# 019-19741), anti-GFAP antibody (mouse, 1:500, Millipore Sigma, Cat# G3893), anti-MBP antibody (Rabbit, 1:500, Boster Bio, Cat# PA1050), and anti-Olig2 antibody (Goat, 1:500, R&D systems, Cat# AF2418). After washing, the sections were incubated with species-specific secondary antibodies conjugated to 488-nm, 555-nm or 633-nm fluorophores (1:500, Jackson ImmunoResearch) for 2 hours at room temperature. Sections were subsequently washed and coverslipped using Fluoroshield™ with DAPI (Sigma, Cat# F6057), and stored in the dark at 4 °C.

### Image acquisition and analysis

After *in situ* hybridization and immunohischemistry, stained sections were imaged using a Zeiss 880 confocal microscope, employing Z-stack imaging. Maximum projections and stitching were accomplished using Zeiss Zen software. Image analysis of percent area coverage of positive staining in the sensory cortex and hippocampus was conducted using Image J software. For each analysis, two to four sections from each mouse were examined, with four mice analyzed per group. All images in a single experiment were captured under the same settings. All analyses and quantifications were performed blinded to the experimental conditions.

### RNA sequencing

Total RNA was extracted from brain tissue (including cortex and hippocampus) of mice with sham surgery, TBI surgery treated with vehicle, and TBI surgery treated with NPD1 mice and processed with RNeasy® Mini Kit (QIAGEN, Cat# 74104). RNA quantity and quality were assessed with a NanoDrop™ One spectrophotometer (Thermo Fisher Scientific). All samples displayed a 260:280 ratio greater than 2.0 and RNA Integrity Numbers (RINs) above 8.0. The RNA concentration > 150 ng/nL and the total RNA yield was > 3 μg. The Poly(A) RNA sequencing library was prepared according to Illumina’s TruSeq-stranded-mRNA sample preparation protocol. RNA integrity was verified using the Agilent Technologies 2100 Bioanalyzer. Poly(A) tail-containing mRNAs were purified using oligo-(dT) magnetic beads, undergoing two rounds of purification. After purification, the poly(A) RNA was fragmented in a divalent cation buffer at elevated temperature, and the DNA library was then constructed. Quality control and quantification of the sequencing library were conducted using Agilent Technologies 2100 Bioanalyzer High Sensitivity DNA Chip. Paired-end sequencing was carried out on the Illumina NovaSeq 6000 sequencing system.

### Bioinformatics analysis

Cutadapt and in-house Perl scripts were utilized to remove adaptor contamination, low-quality bases, and undetermined bases from the reads. Subsequently, sequence quality was verified using FastQC (http://www.bioinformatics.babraham.ac.uk/projects/fastqc/). Reads were mapped to the mouse genome using HISAT2 (ftp://ftp.ensembl.org/pub/release-101/fasta/mus_musculus/dna/). The mapped reads from each sample were assembled using StringTie. All transcriptomes were then merged to reconstruct a comprehensive transcriptome, employing Perl scripts and gffcompare. Once the final transcriptome was generated, StringTie and Ballgown were used to estimate the expression levels of all transcripts. mRNA expression levels were quantified by calculating FPKM values using StringTie. Differential expression analysis of mRNAs was performed using the R packagelimma and edgeR. A total of 35,797 genes were recorded and 21,414 genes were further analyzed. Analysis was performed using R studio version 2023.09.1+494 (R version 4.3.2) according to Limma and edgeR packages workflow(*51*). Briefly, original counts from each sample were integrated into a DGEList object and low expressed genes were filtered out by filterByExpr method from edgeR package. Normalization was conducted by the Trimmed Mean of M-values (TMM) method. Genes with the parameter of false discovery rate (FDR) below 0.05 and absolute fold change ≥ 2 were defined as differentially expressed genes. Venn diagram was made by venn.diagram method from VennDiagram package. ENTREZ IDs were matched with gene symbols using the org.Mm.eg.db package. GO analysis was performed with enrichGO method from clusterProfiler package.

### Computational simulation

Molecular modeling and simulation analysis was performed as previously described(*27*). The structure of NPD1 was retrieved from TargetMol. Molecular dynamics simulations (MDS) were conducted using DESMOND software in Schrödinger Release 2018-4(*52*). The stability of the protein-ligand complex was assessed through MDS, with trajectory clustering to analyze conformational changes. Based on atomic root mean square deviation (RMSD), 1000 trajectory structures were clustered into ten clusters using Desmond’s trajectory clustering method. Finally, 1000 ns MDS was conducted on the top three representative clusters to evaluate the stability of the docking complex.

### Quantification and statistical analysis

All data were expressed as the mean ± SEM. Number of experimental replicates (n) is indicated in the figure legends and refers to the number of experimental subjects independently treated in each experimental condition. The criterion for statistical significance was defined as *P* < 0.05. Asterisks indicate the following significance levels: \**P* < 0.05, \*\**P* < 0.01, \*\*\**P* < 0.001, \*\*\*\**P* < 0.0001. Statistical analyses were completed with Prism GraphPad 10.0. All the experiments in this study were done and analyzed by researchers who were blinded to genotype or treatment.

## Acknowledgements

This study was supported by DoD grants W81XWH2110885 and W81XWH2110756 and NIH R01 grant NS131812. The Schematic figures are prepared with BioRender with license.

## Author contributions

J.L.Z. and R.R.J designed the project, interpreted the data and wrote the paper with input from all authors. J.L.Z., H.C.W performed the experiments, S.C performed the computer stimulation, Y.Q.W analyzed the RNAseq data. V.Z. edited the manuscript with critical comments. All authors discussed and edited the manuscript; all authors approved the final manuscript submission.

---

**Fig. S1.**
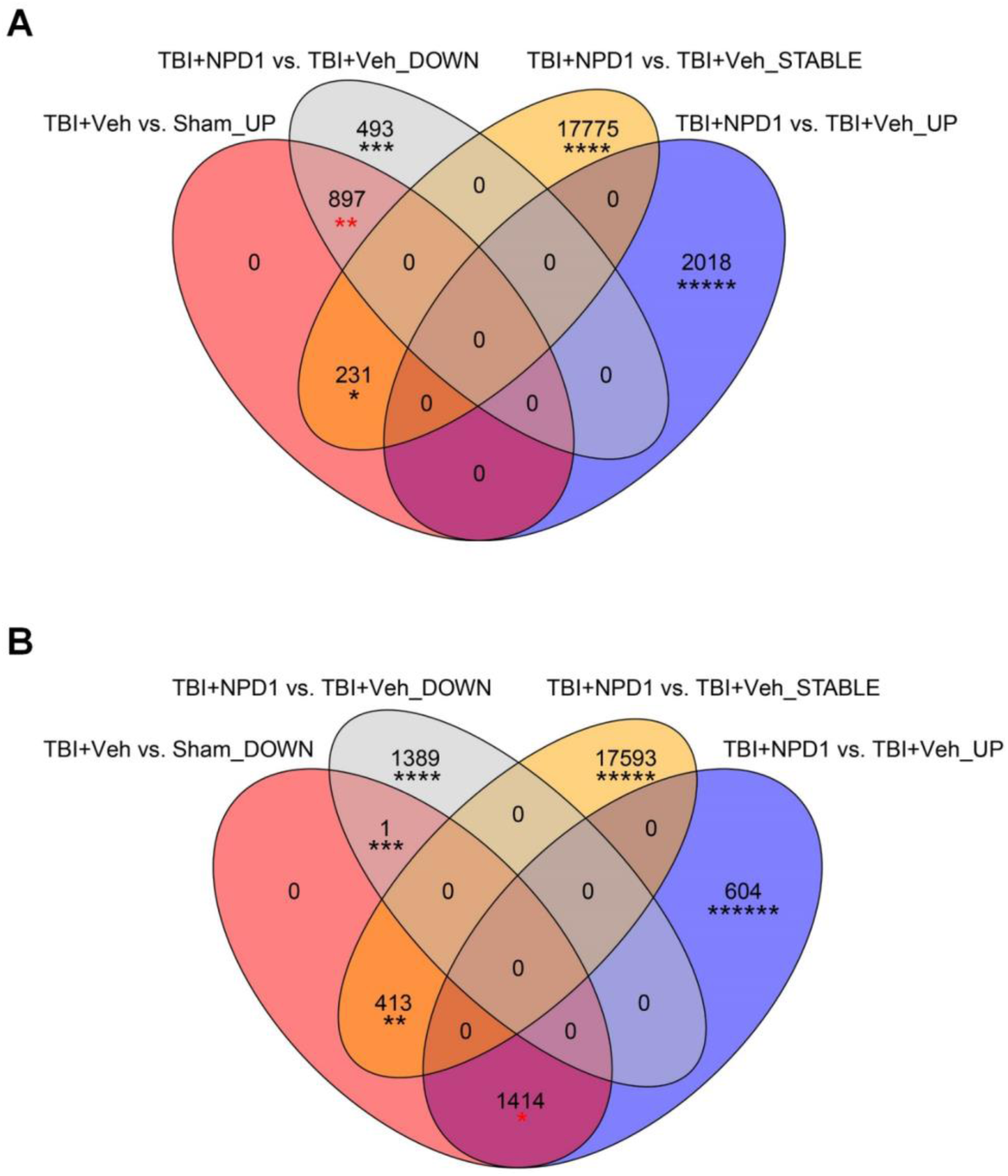
Numbers of overlapping differentially expressed genes in mice with sham surgery, TBI treated with vehicle, and TBI treated with NPD1. **(A)** The overlapping differentially expressed genes which are upregulated by TBI but are downregulated by NPD1 treatment. * indicates 231 genes that are upregulated by TBI Surgery (vs. Sham) but not restored by NPD1 treatment. ** indicates 897 genes that are upregulated by TBI (vs. Sham) and further downregulated by NPD1 treatment; *** indicates 497 genes that are only downregulated by NPD1 treatment (vs TBI); **** indicates 17,775 genes that are stable after TBI surgery and NPD1 treatment; ***** indicates 2,018 genes that are only upregulated by NPD1 treatment (vs. TBI). **(B)** The overlapping differentially expressed genes which are downregulated by TBI and upregulated by NPD1 treatment. * indicates 1,414 genes that are downregulated by TBI (vs. Sham) but are also upregulated by NPD1 treatment; ** indicates 414 genes that are downregulated by TBI but cannot be restored by NPD1 treatment; *** indicates only 1 gene that is downregulated by both TBI surgery and NPD1 treatment; ****indicates 1,389 genes that are only downregulated by NPD1 treatment (vs. TBI); ***** indicates 17,593 genes that are stable after TBI surgery and NPD1 treatment; ****** indicates 604 genes that are downregulated by NPD1 treatment (vs. TBI). Data were collected from three groups of mice, including sham surgery (Group-1), TBI surgery treated with vehicle (Group-2) and TBI surgery treated with NPD1 (Group-3). Note that NPD1 treatment can substantially normalize the dysregulated genes by TBI.

**Fig. S2.**
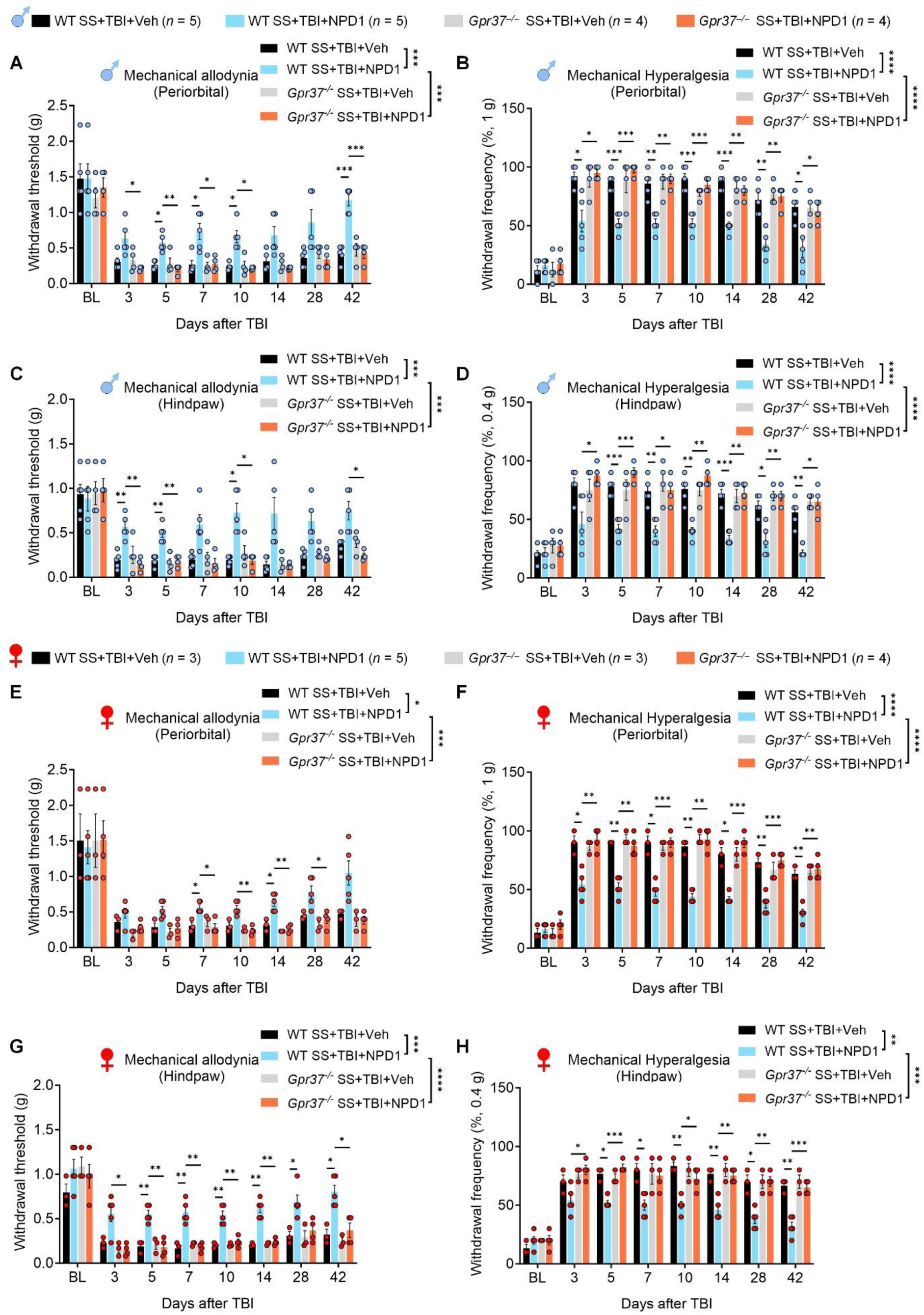
NPD1 is effective in preventing stress-induced chronification of pain in both sexes of TBI mice with swimming stress. **(A-D)** Primary and secondary mechanical pain in male mice. **(A, B)** Primary mechanical pain: von Frey testing showing withdrawal threshold (A) and withdrawal frequency (B) at periorbital site in four groups. **(C, D)** Secondary mechanical pain: von Frey testing showing withdrawal threshold (C) and withdrawal frequency (D) at hindpaw site in four groups. **(E-H)** Primary and secondary mechanical pain in female mice. **(E, F)** Primary mechanical pain: von Frey testing showing withdrawal threshold (E) and withdrawal frequency (F) at periorbital site infour groups. **(G, H)** Secondary mechanical pain: von Frey testing showing withdrawal threshold (G) and withdrawal frequency (H) at hindpaw site in four groups. Group 1: Swimming stress (SS), TBI surgery and behavioral test in WT mice with vehicle treatment. Group 2: SS, TBI surgery and behavioral test in WT mice with NPD1 treatment. Group 3: SS, TBI surgery and behavioral test in *Gpr37^-/-^* mice with vehicle treatment. Group 4: SS, TBI surgery and behavioral test in *Gpr37^-/-^* mice with NPD1 treatment. Also see Fig. 6a. Data are represented as mean ± SEM. \**P* < 0.05, \*\**P* < 0.01, \*\*\**P* < 0.001, \*\*\*\**P* < 0.0001. Two-way ANOVA testing for two group comparison (A-H), Two-way ANOVA followed by Tukey’s multiple comparisons test for time point comparison (A-H).

**Fig. S3.**
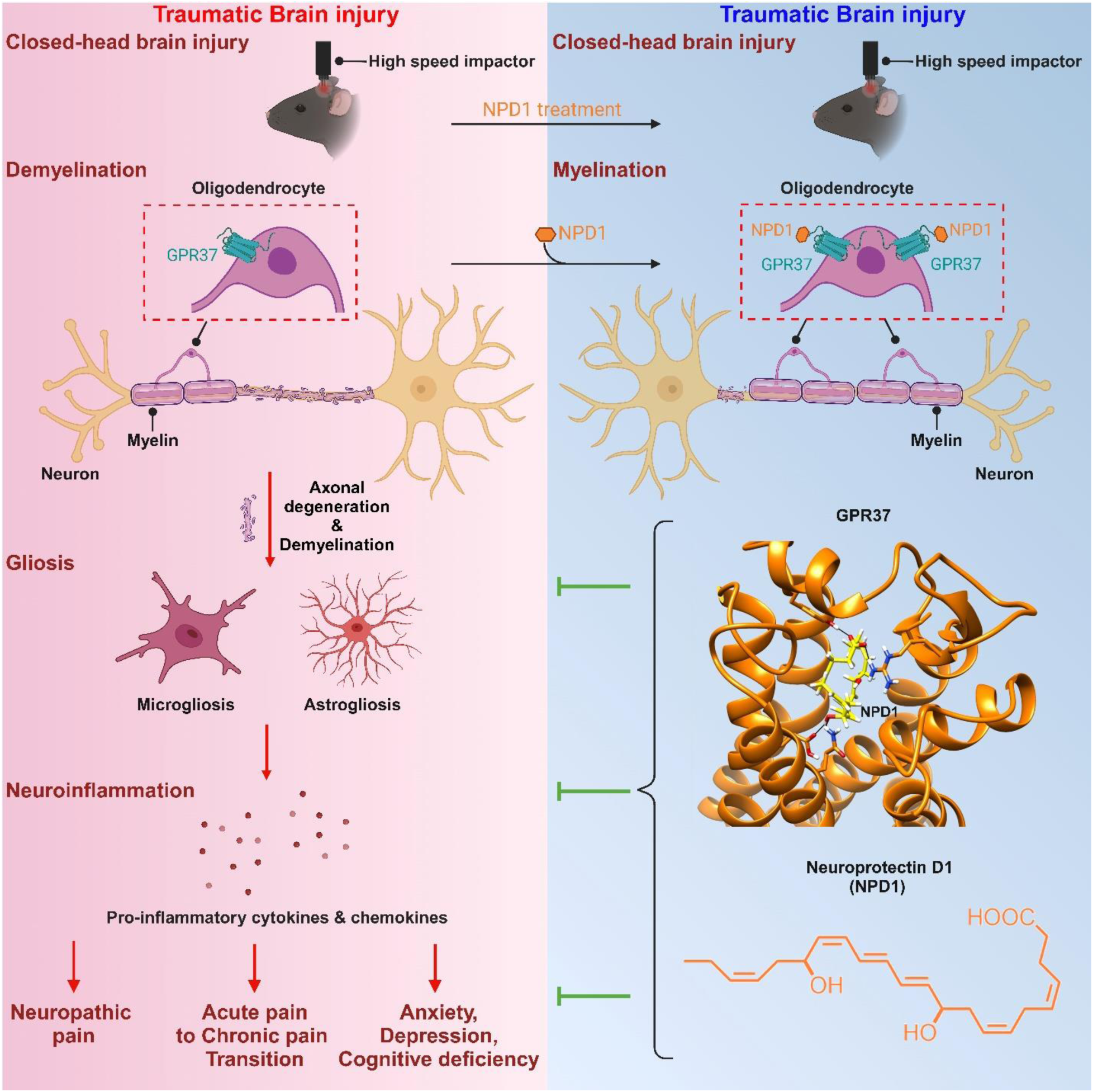
Schematic of the working hypothesis for the protective effect of NPD1/GPR37 signaling pathway in TBI induced demyelination, gliosis, neuroinflammation as well as neuropathic pain and its complications.

